# Common Piezo1 allele in African populations causes xerocytosis and attenuates Plasmodium infection

**DOI:** 10.1101/159830

**Authors:** Shang Ma, Stuart Cahalan, Rakhee Lohia, Gregory LaMonte, Weizheng Zeng, Swetha Murthy, Emma Paytas, Nathan D. Grubaugh, Ramya Gamini, Laurence Berry, Viktor Lukacs, Tess Whitwam, Meaghan Loud, Andrew I. Su, Kristian G. Andersen, Elizabeth A. Winzeler, Eric Honore, Kai Wengelnik, Ardem Patapoutian

**Affiliations:** Howard Hughes Medical Institute, Molecular and Cellular Neuroscience, Dorris Neuroscience Center, The Scripps Research Institute, La Jolla, CA 92037, USA.; DIMNP, CNRS, INSERM, Univ Montpellier, Montpellier, France.; Division of Host-Microbe Systems & Therapeutics, Department of Pediatrics, University of California, San Diego, California, USA.; Department of Immunology and Microbiology, The Scripps Research Institute, La Jolla, California, USA.; Department of Integrative, Structural and Computational Biology, Scripps Research Institute, La Jolla, California, USA.; Université Côte d’Azur, CENTRE NATIONAL DE LA RECHERCHE SCIENTIFIQUE ^+^Institut de Pharmacologie Moléculaire et Cellulaire, Valbonne, France

## Abstract

Hereditary xerocytosis (HX) is thought to be a rare genetic condition characterized by red blood cell (RBC) dehydration with mild hemolysis. Gain-of-function (GOF) mutations in mechanosensitive Piezo1 ion channels are identified in HX patients. RBC dehydration is linked to reduced *Plasmodium* infection rates in vitro. We engineered a Piezo1 mouse model of HX and show that *Plasmodium* infection fails to cause experimental cerebral malaria in these mice. Furthermore, we identified a novel GOF human Piezo1 variant, E756del, present in a third of African population. Remarkably, RBCs from individuals carrying this allele are dehydrated and protected against *Plasmodium* infection in vitro. The presence of an HX-causing Piezo1 mutation at such high frequencies in African population is surprising, and suggests an association with malaria resistance.

---

*Plasmodium*, the causative parasites for malaria, has exerted strong selective pressure on the human genome. This is demonstrated by severe genetic conditions, such as sickle cell disease (SCD), that persist in certain populations because they confer resistance to *Plasmodium* infection (1). The high frequency of genetic mutations that causes sickled red blood cells (RBCs) in malaria-endemic regions suggests a balancing selection for SCD and malarial resistance that still continues in present days (2, 3, 4). However, the scope of RBC disorders that might contribute to *Plasmodium* resistance has not been fully explored. Hereditary xerocytosis (HX) is thought to be an extremely rare condition almost exclusively found in Caucasian families (5). It is characterized by RBC dehydration and can cause mild anemia or, more often, be asymptomatic. Recently, several gain-of-function (GOF) mutations in Piezo1 were identified in HX patients (6, 7, 8) and shown to be associated with this genetic disorder. Piezo1 is a mechanically activated ion channel and is shown to control RBC volume regulation (9). Interestingly, dehydrated RBCs (including those from HX patients) are somewhat resistant to *P. falciparum* infection in vitro (10).

To confirm that Piezo1 GOF causes HX and to test the role of HX in malaria infection in vivo, we engineered a mouse model in which we conditionally expressed a human-equivalent HX mutation (Fig.1). Specifically, R2456H is one of the mutations in human Piezo1 that displays significantly slower inactivation [i.e. longer channel inactivation time (τ)], is considered a gain-of-function (GOF) mutation, and is associated with HX in linkage analyses (5). We first verified that the equivalent mouse Piezo1 (mPiezo1) point mutation (R2482H), when overexpressed in HEK cells that lack endogenous Piezo1 (Piezo1 KO HEK) (11), also shows a similar shift in the inactivation kinetics of mechanically activated currents (Fig.1A). Since residue 2482 resides in the last coding exon, we designed the knock-in construct by flanking exons 45-51 with flox sites, followed by a copy of the exons 45-51 region with a mutation that would replace R to H at residue 2482 (Fig.1B). When Cre recombinase is expressed, the wild type exon will be replaced by the modified exon, enabling tissue-specific Piezo1 GOF expression. To characterize the knock-in mouse model, we generated a blood cell-specific Piezo1 GOF mouse line by crossing Piezo1cx/cx into vav1-cre, a Cre line that targets all hematopoietic lineages (12). We sequenced the last exon of piezo1 cDNA from Piezo1cx/cx; vav1-cre blood, and observed the expected nucleotide change (Fig. S1A). In addition, we found that the piezo1 transcript levels in whole blood from both Piezo1cx/cx; vav1-cre and Piezo1cx/+; vav1-cre mice were not different from wild type, demonstrating that the genetic manipulation does not alter Piezo1 expression levels (Fig. S1B). The knock-in mice were born at the expected Mendelian ratio and appear to develop normally. Importantly, red blood cells from both Piezo1cx/cx; vav1-cre and Piezo1cx/+; vav1-cre showed reduced osmotic fragility, as shown by a left shifted curve in a hypotonicity-dependent hemolysis challenge (Fig. 1C-D). This suggested that RBCs from Piezo1 GOF knock-in mice are more resistant to lysis in response to hypotonic solutions compared to wild type, a defining feature for HX (13). In addition, we assessed the hematological properties of whole blood from Piezo1 GOF mice (Table S1). As in HX patients, both heterozygote and homozygote knock-in mice display mild anemia, indicated by a lower hemoglobin (HGB) level and increased reticulocyte number (RET). Shifts in Mean Corpuscular Volume (MCV) and Mean Corpuscular Hemoglobin (MCH) values are reminiscent of HX patients. In addition, scanning electron microscopy showed that some RBCs from piezo1cx/+; vav1-cre have deformed and dehydrated shapes, as is often observed in blood from HX patients (Fig. 1E). Spleen is the major site for removing abnormal RBCs during blood circulation, and HX patients can experience splenomegaly. Indeed, both Piezo1cx/cx; vav1-cre and Piezo1cx/+; vav1-cre mice have significantly larger spleens compared to wild type mice (Fig. 1F and Fig. S1C). Therefore, our blood-specific Piezo1 GOF model recapitulates HX conditions and it provides a unique in vivo setting to study this inherited disease, as well as explore the connection between RBC dehydration and blood-borne parasitic diseases in vivo.

**Figure 1.**
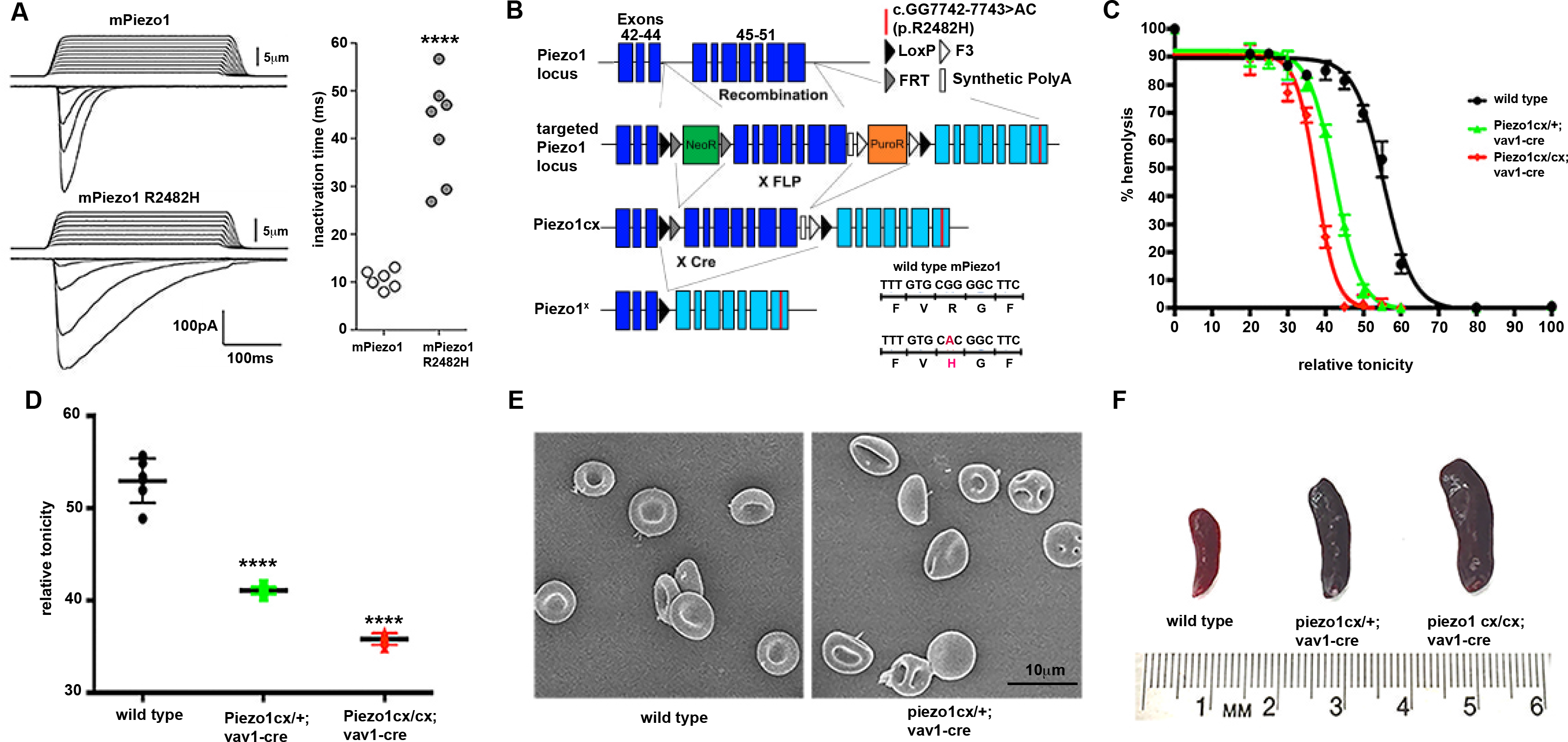
Mouse model for human xerocytosis. (A) Representative traces of mechanically-activated (MA) inward currents in Piezo1 KO HEK cells expressing wild type and Piezo1 R2482H. mPiezo1 R2482H mutations had normal MA currents but showed significantly longer inactivation time (****p < 0.001, right panel). (B) Strategy for generating knock-in mouse. Recombinase activity removes endogenous wild type exons, only expressing the modified exon that has the R2482H mutation. (C) Osmotic fragility test for red blood cells (RBCs). A left-shifted curve (green and red) compared to wild type (black) means reduced fragility, which indicates RBC dehydration. (D) Quantification for osmotic fragility. Relative tonicity at which 50% RBCs are lysed (half hemolysis) was calculated for each curve. Piezo1cx/cx; vav1-cre and Piezo1cx/+; vav1-cre mice have significantly reduced osmotic fragility than wild type (****p < 0.001). (E) Scanning electron microcopy images. Piezo1cx/+; vav1-cre RBCs showed abnormal shapes indicating stomatocytes similar to human HX patients. (F) Splenomegaly in Piezo1 GOF mice. Spleens from Piezo1 cx/+; vav1-cre and Piezo1 cx/cx; vav1-cre are significantly larger than wild type. (Fig S1C).

Next, we infected the knock-in mice with a rodent specific *Plasmodium* species, *P. berghei* ANKA, engineered to express GFP to enable fluorescence-based parasitemia analysis (14). Intravenous injection of 1x10^5^ parasite-infected RBCs caused wild type C57B/6 mice to die between day 6 and 8, consistent with previous literature (15) (Fig. 2A). To evaluate the function of Piezo1 GOF in malarial infection, we generated constitutive knock-in Piezo1 GOF mice by crossing Piezo1cx/cx with cmv-cre, a cre driver expressed ubiquitously (16). Remarkably, Piezo1 cx/cx; cmv-cre and Piezo1 cx/+; cmv-cre mice (all on C57B/6 genetic background, described in Methods) survived as long as 24 and 19 days, respectively, suggesting that Piezo1 GOF confers protection against *Plasmodium* infection in vivo (Fig. 2A). Importantly, the blood cell-specific Piezo1 GOF mice, both Piezo1cx/cx; vav1-cre and Piezo1cx/+; vav1-cre, had post-infection survival rates indistinguishable from constitutive knock-in animals (for both heterozygotes and homozygotes), suggesting that Piezo1 GOF in hematopoietic lineages is sufficient to drive the protection (Fig. 2A).

**Figure 2.**
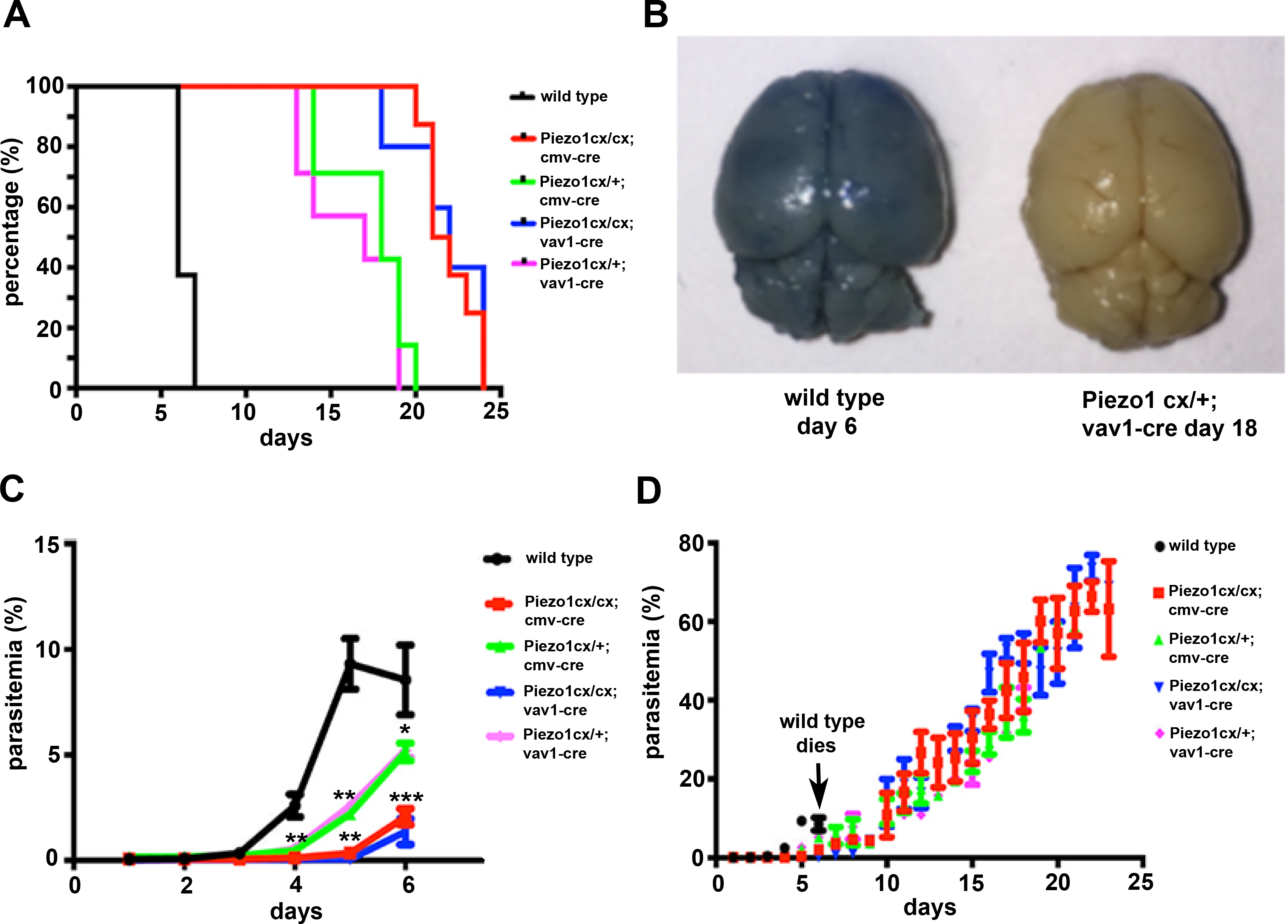
Protection against *Plasmodium* by Piezo1 gain-of-function. (A) Kaplan-Meier survival curves for Piezo1 GOF mice after intravenous injection of 1×10^5^ *P. berghei* infected RBCs (iRBCs). (B) Intact blood brain barrier (BBB) in infected Piezo1 GOF mice. Brains were dissected for analysis after Evan blue injection from morbid mice that showed hunching, ataxia (wild type) or immobility (knock-in). Piezo1cx/+; vav1-cre did not show compromised BBB. (C) and (D) Parasitemia recorded by flow cytometry for 6 days and 24 days respectively.

Cerebral malaria is the most pernicious form of human severe malaria, causing more acute deaths than any other malarial symptoms including severe anemia (17). To evaluate the occurrence of cerebral malaria in our model, Evans blue dye was injected into infected mice. This showed blue dye leakage into brain parenchyma on day 6 in all wild type mice (n=8) due to a disrupted blood-brain barrier (BBB) (Fig. 2B, wild type), indicating experimental cerebral malaria (ECM). Remarkably, the BBB of Piezo1cx/+; vav1-cre mice (n=7) remained intact even at the end of their life, suggesting that Piezol GOF expression completely protects animals from BBB breakdown, a key component of ECM (Fig. 2B). Next, we analyzed the parasitemia post *P. berghei* infection in wild type and knock-in animals by measuring the percentage of RBCs that are GFP-positive by flow cytometry. During the first week of infection, knock-in animals (both constitutive and blood-specific) had significantly lower parasitemia compared to wild type, suggesting that expression of Piezo1 GOF allele in blood cells reduces *Plasmodium* development in vivo (Fig. 2C). Parasitemia reached 6-12% in wild type animals at time of death (Fig. 2C). Knock-in animals exhibited a slow increase in parasitemia, eventually leading to severe hyperparasitemia (up to 70% parasitemia), but did not develop signs for ECM (Fig. 2D). Eventually, Piezo1 GOF mice succumbed to severe anemia instead of cerebral malaria, since the HGB level was significantly lower (2.85±2.5 g/dL, n=3) in Piezo1 cx/+; vav1-cre blood 18 days after infection compared to uninfected knock-in mice (14.02± 0.16 g/dL, n=5, p<0.002) (18). The decrease in parasitemia observed in knock-in mice during the first week is readily explained by reduced infection rates due to dehydrated RBCs (see also below, Fig. 3). The lack of ECM that leads to hyperparasitemia is potentially due to altered immune responses to *P. berghei infection* in Piezo1 GOF mice. This may arise due to cell autonomous roles of Piezo1 GOF in white blood cells.

**Figure 3.**
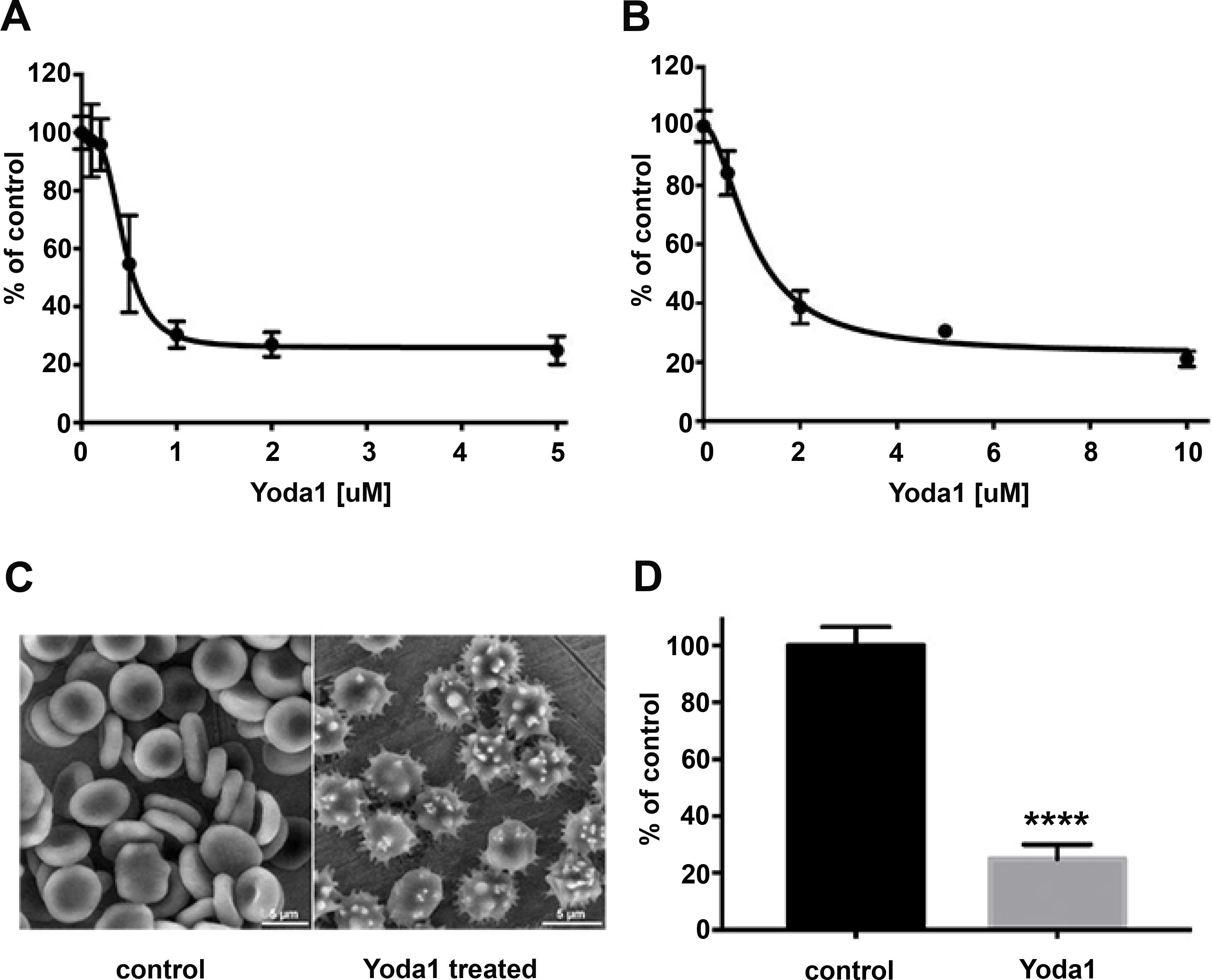
*P. falciparum* infection of human RBCs in response to Yoda1. (A) Effect on parasitemia in the presence of Yoda1 over one full developmental cycle. Treatment was started on highly synchronized ring stage parasite cultures with Yoda1 containing medium change every 24 h. Final parasitemia was determined after 60 h. (B) Effect of Yoda1 treatment during the transition from schizonts to ring stages. Highly synchronized schizont cultures were treated and parasitemia determined after 18 h. (C) RBC shape change induced by Yoda1. In the presence of 15 pM Yoda1 (right panel), human RBCs showed echinocyte morphology in scanning electron microscopy in comparison to the biconcave disk shape of untreated controls (left panel). (D) Pre-treatment of RBC with Yoda1 inhibits parasite development. Uninfected RBC were treated for 10 min with 15 μM Yoda1, washed thrice and mixed with untreated purified late schizont infected RBCs. Parasitemia was determined after 18 h. Data for A-C were expressed as percentage of mock treated controls, ****p<0.001, Student’s t-test

Alternatively, slower parasite replication may allow Piezo1 GOF mice to develop increased ability to mount a protective immune response. To test if this protection was an indirect effect secondary to initial low infection rates (Fig. 2C), we infected wild type mice with reduced pathogen load (4×10^4^ parasitized RBCs) and showed that reduced initial infection rate does not by itself give rise to the phenotype observed in knock-in mice (Fig. S2A). Together, our data demonstrate that overactive Piezo1 in blood cells considerably reduces *Plasmodium* growth and protects mice from cerebral malaria. We hypothesize that RBC dehydration (also associated with HX) is a leading cause of the reduced parasitemia observed in Piezo1 GOF animals. However, the mechanism involved in protection from severe malaria will have to await future investigations into a potential role of GOF Piezo1 allele in RBCs and/or in immune cells.

The genetic data above suggest that RBC dehydration plays an important role in reducing *P. berghei* infection during the first week in vivo. To explore whether acute RBC dehydration induced by Piezo1 activation is also protective in human RBCs against *P. falciparum* infections, we used Yoda1, the Piezo1-specific agonist (19) to treat human RBCs and then infected these cells with *P. falciparum*, in vitro. Culturing human RBCs with *P. falciparum* in the presence of Yoda1 dose-dependently reduced the parasitemia after 48h by up to 80% (Fig 3A). A strong reduction was already observed when purified late stage parasites were incubated in the presence of Yoda1 and parasitemia was determined 18h later at the ring stage (Fig 3B). Scanning electron microscopy showed that Yoda1 treated RBCs displayed a morphology resembling human echinocytes, with a typical dehydrated shape (Fig 3C). To confirm that the effect of Yoda1 is through activation of RBC Piezo1 and not through direct action on the parasite (even though *Plasmodium* species do not have Piezo orthologues), human RBCs were first pre-incubated with Yoda1 for 10 min, extensively washed and then mixed with purified late stage parasites that were close to egress. Again, next day ring stage parasitemia was strongly reduced with respect to controls, revealing that the reduction in parasitemia is due to an overactive Piezo1 channel and not to Yoda1 activity on the parasite (Fig 3D). It also indicated that the short Yoda1 pre-treatment had a protective effect for a prolonged period of time. Taken together, these observations confirm that the results we observed in mice are also relevant to human RBCs (10).

These findings raise a conundrum: If HX is protective against malaria infection and cerebral malaria, why is then HX not described at all in people of African descent, where malaria is highly prevalent? Given the mild phenotype of HX patients (often asymptomatic) (13), one could expect selection of Piezo1 mutations that cause HX in areas where malaria is endemic. One possible explanation is that xerocytosis is often asymptomatic and is likely to be overlooked, especially in understudied African populations. Therefore, we took a comparative genomics approach to look for possible Piezo1 GOF alleles in African populations. We were interested in missense single nucleotide polymorphisms (SNP) and in-frame insertions/deletions (indels) of Piezo1 mutations with reasonably high allelic frequencies (>0.5%) that would be more prevalent in African than non-African populations (>5-fold enriched compared to European population). Analysis of the Exome Aggregation Consortium data (ExAC) (20) revealed 21 mutations that satisfied the above criteria (Table S2). We next expressed full-length human Piezo1 cDNAs containing each of the 21 mutations into Piezo1KO HEK cells, and performed an intracellular calcium assay to screen for GOF mutations. We identified two mutations (amino acid substitution A1988V and deletion E756del), that when overexpressed, showed increased calcium signals compared to the wild type Piezo1 in response to different concentrations of Yoda1 (Fig 4A-B). A1988V mutation only exists in African population with 0.8% allelic frequency (inset in Fig. 4A). Surprisingly, we found that E756del has an allelic frequency of 18% in African population (inset in Fig 4B), such that approximately one third of the African populations carry at least one copy of the E756del mutation (Fig 4E). To confirm if these mutations are indeed GOF mutations, we recorded mechanically activated currents and found that both A1988V and E756del mutations were activated normally by mechanical force but had significantly longer inactivation times (τ) compared to wild type, similar to R2456H, a GOF allele with the longest inactivation time among all HX mutations (6) (Fig 4C-D). Therefore, Piezo1 GOF mutations, similar to those causing HX in Caucasian families (6), exist in African population. We used population genetics to determine if A1988V and E756del display signatures of being under positive selection in African populations. While the mutation A1988V was only detected in African populations, its frequency remains low (0.8%, inset Fig. 4A) and is unlikely to be strongly selected (Table 2S). E756del, however, has high allelic frequencies (on average 18%) in all African populations (inset in Fig B and 4E), and a third of sampled individuals of African descent carry at least one copy of the deletion (Figure 5). The observed genotype frequencies at this locus are as expected for random mating and in Hardy-Weinberg equilibrium (*χ* ^2^ =0.201, p = 0.654). Since we did not observe any other SNPs in linkage disequilibrium with E756del in our analysis, it is difficult to directly assess signatures of selection around this variant. However, E756del has the highest degree of variance between the African and non-African populations compared to all other Piezo1 missense alleles from the 1000 Genomes database (21) (F_ST_ for E756del = 0.32, F_ST_ of all other Piezo1 missense alleles = 0-0.26, Table 2S and Fig 3S). These findings are consistent with positive selection for E756del in African populations, however, do not provide conclusive evidence (22).

**Figure 4.**
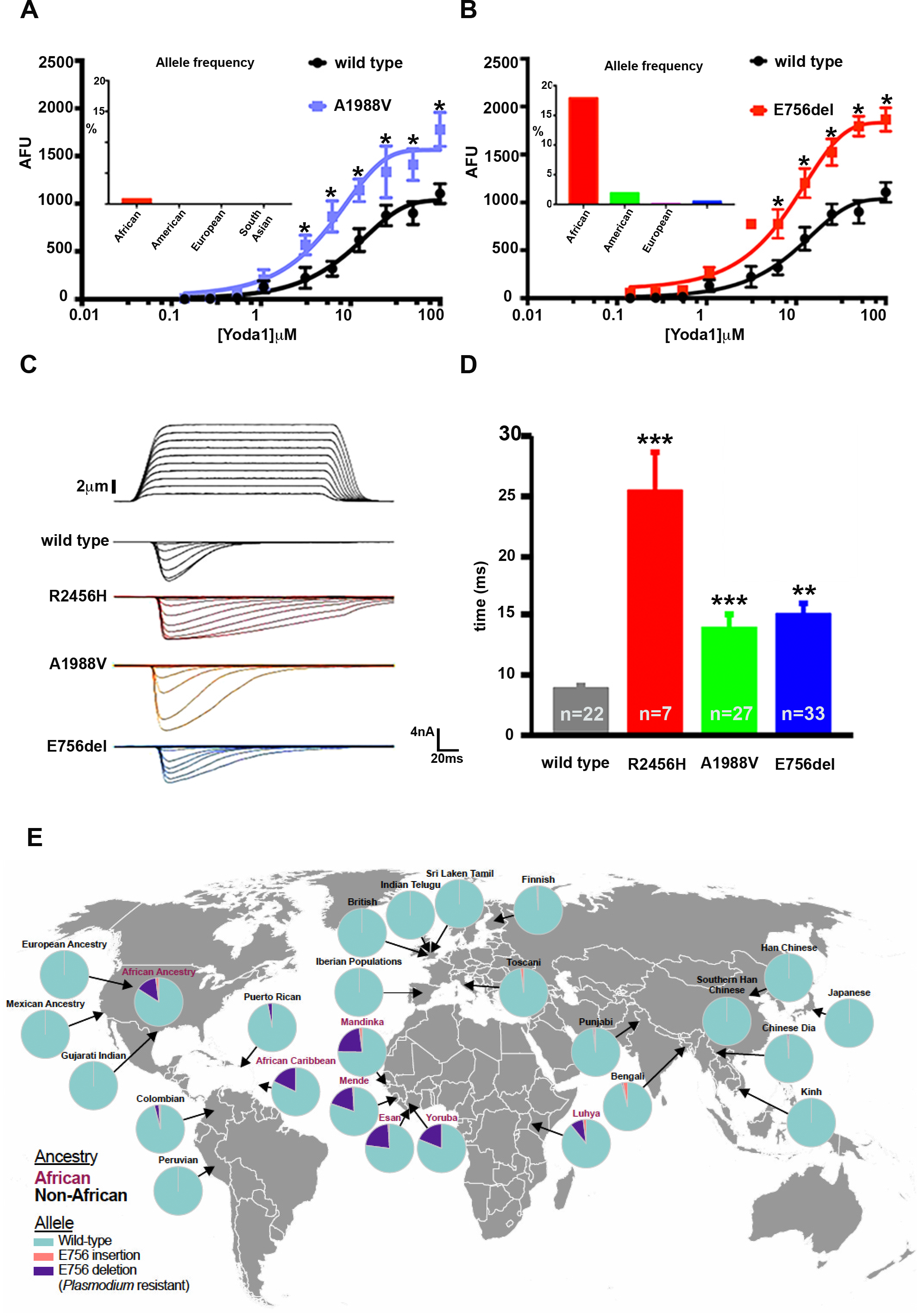
Identification of Piezol gain-of-function mutations in African populations. (A) and (B) Yodal induced Intracellular calcium signals in PiezolKO HEK cells overexpressing A1988V and E756del cDNA. A1988V and E756del have significantly higher signals than wild type (* p<0.05). Allele frequency for both mutations is shown in the insets. (C) Representative traces of mechanically activated (MA) inward currents in Piezo1 KO HEK cells expressing wild type and mutated cDNA. R2456H (human xerocytosis mutation), A1988V, and E756del mutations had normal MA currents similar to wild type but they showed significantly slower inactivation kinetics. (D) Quantification for inactivation time (T). n is the number of cells. (***p < 0.001, **p < 0.01). (E) Human population demographics for E756 indels. E756del exists at high frequencies in all populations of African descents (purple).

**Figure 5.**
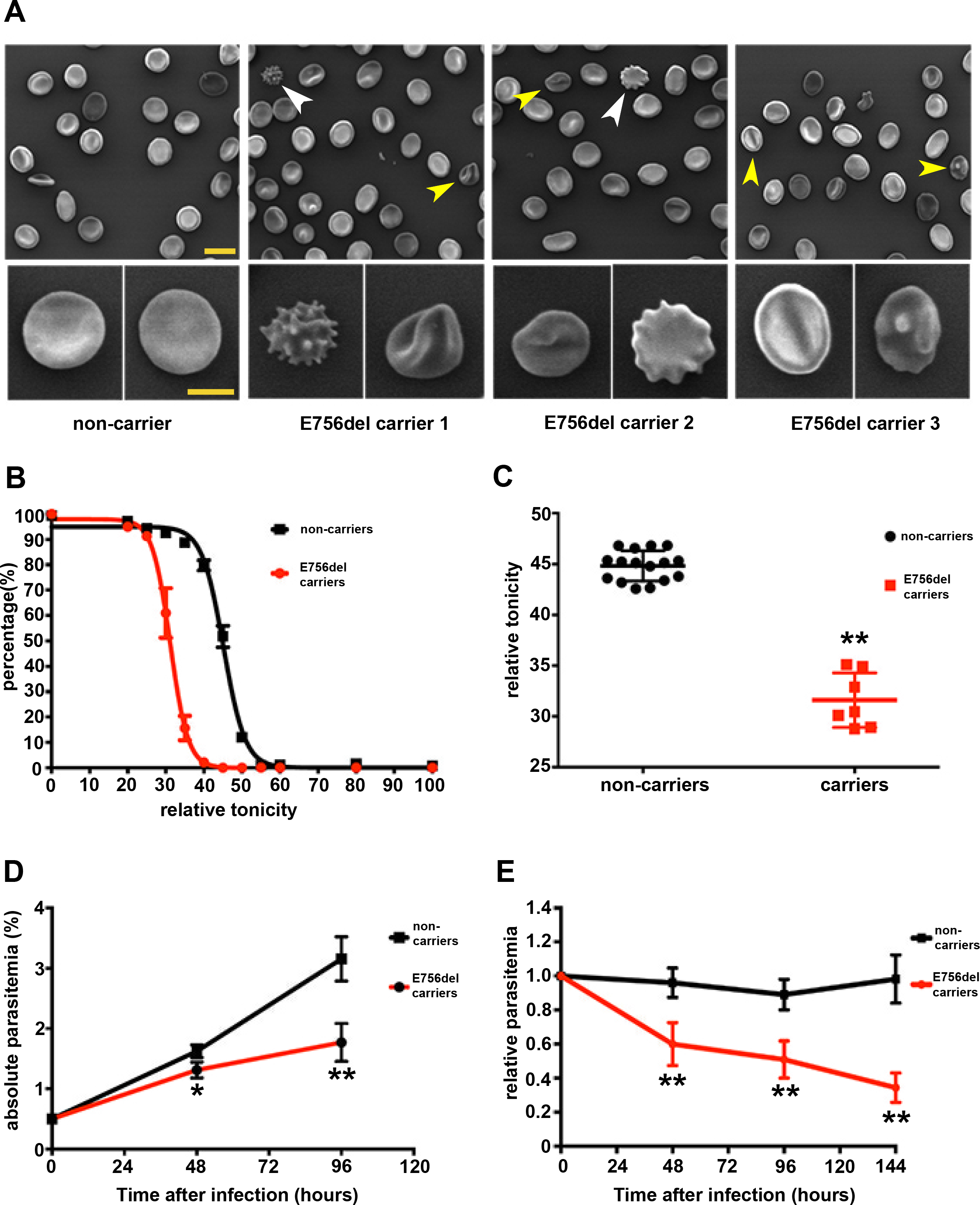
Characterization of RBCs from E756del carriers for xerocytosis-like phenotypes, and *P. falciparum* infection. (A) SEM images. Three individual E756del carriers have RBCs with echinocytes (white arrowhead) and stomatocytes (yellow arrowhead), also magnified in lower panels. Scale bar for upper panels: 10um; for lower panels: 5um. (B) and (C) Osmotic fragility test. RBCs from E756del heterozygous carriers (n=9) had a left-shifted curve (red) compared to the non-carrier control (n=16) (black), indicating a reduced fragility, as quantified in (C) **p <0.01. (D) and (E) *P. falciparum* infection into RBCs from E756del carriers. Giemsa staining (D) and SYBR Green labeling of parasite DNA inside RBCs (E) showed that parasitemia for RBCs is significantly lower for E756del carrier (n=9) compared to wild type (n=16) (**p < 0.01, *p < 0.05). Statistics: student’s t-test for each time point.

Because of its high frequency (in contrast to the A1988V), we were able to acquire blood samples from healthy volunteer African American donors to test whether E756del causes HX-like RBC dehydration, and importantly, whether it confers resistance to *P. falciparum* infections of RBCs. We obtained 25 whole blood samples and used white blood cells to sequence the exon containing E756del. Nine of the donors were tested heterozygote for E756del. Scanning electron microscopy images showed the presence of RBCs with echinocyte and stomatocyte morphologies in all three carriers we examined (Figure 5A). Remarkably, we found that RBCs from all 9 donors were dehydrated (Fig. 5B-C), also similar to RBCs from known xerocytosis patients. This suggests that RBC dehydration (due to expression of Piezo1 GOF allele) is unexpectedly common in African populations. Next, we infected both non-carrier and E756del carrier RBCs with *P. falciparum* in vitro (genotypes blinded from experimenters) (23). Our data showed that parasitemia was significantly lower for E756del heterozygous carriers relative to non-carriers at both 48hr and 96hr post infection (Fig. 5D). In addition, we used a more quantitative method based on SYBR green labeling of parasite DNA to measure the parasitemia. Our results show that parasitemia (normalized to the non-carrier control) for carriers’ blood was significantly lower than controls at 48hr, 96hr and 144hr post infection (Fig. 5E), consistent with Giemsa staining data. Together, our data demonstrate that E756del is a common Piezo1 GOF mutation in African populations and that it confers resistance to RBC infection by *P. falciparum* at least in vitro.

Here, we generated a mouse xerocytosis model and demonstrate that overactive mechanically activated Piezo1 channel in blood cells strongly protects animals from *Plasmodium* infection by significantly reducing RBC infection and preventing cerebral malaria. Strikingly, we identified a Piezo1 GOF mutation prevalent in populations of African descent that causes dehydrated RBCs, similar to HX patients. This is surprising, as HX is thought to be a very rare disorder that mainly afflicts Caucasians (6, 7, 8). The donors that carry this allele are apparently asymptomatic as they were registered as healthy blood donors. However, we do not have any health records associated with these individuals, and full clinical evaluation of individuals carrying this allele will be of high interest. Remarkably, the mutation considerably reduces *P. falciparum* infection rates. We conclude that HX-like condition is much more common than previously anticipated and might confer protection against *P. falciparum.* Despite the experimental evidence above, E756del was not detected as a strong candidate by recent genome-wide association studies (GWAS) that aimed to identify genetic loci for severe malaria resistance (24). This is potentially due to GWAS limitations and the complexity of this particular genetic locus. GWAS samples have high levels of genetic diversity and are underrepresented in reference panels of genetic variation (25). Also, E756del is in a region with multiple short tandem repeats so that imputation of this mutation into current GWAS datasets is not straightforward. In the future, analysis of epidemiological data will reveal whether this Piezo1 variant is indeed associated with protection against severe malaria (25). In addition, clinical characterization of individuals with the E756del allele will shed further light on the range of phenotypes that are associated with Piezo1 GOF expression, including anemia, splenomegaly, as well as various aspects of cardiovascular function (26, 27, 28).

---

**Figure S1.**
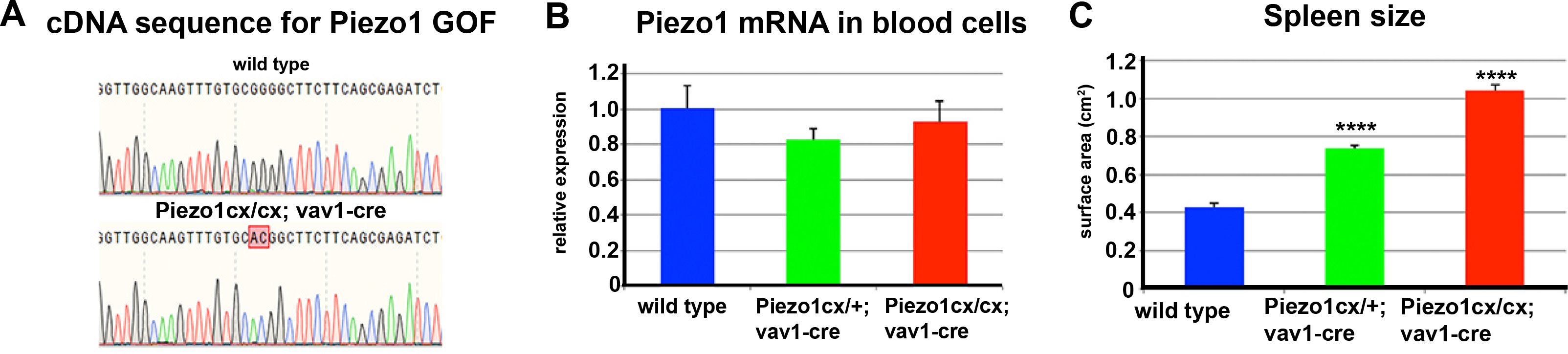
(A) Modified nucleotides in the last exon of piezo1 cDNA from Piezo1cx/cx; vav1-cre (lower). GG to AC change compared to wild type (upper). (B) piezo1 transcript levels in heterozygotes and homozygotes for Piezo1 GOF. p >0.05 (One-way ANOVA test). (C) Spleens from Piezo1 cx/cx; vav1-cre or Piezo1 cx/+; vav1-cre are significantly larger than wild type. ****p <0.0001, student’s t-test, both compared to wild type (n=3 for each group).

**Figure S2.**
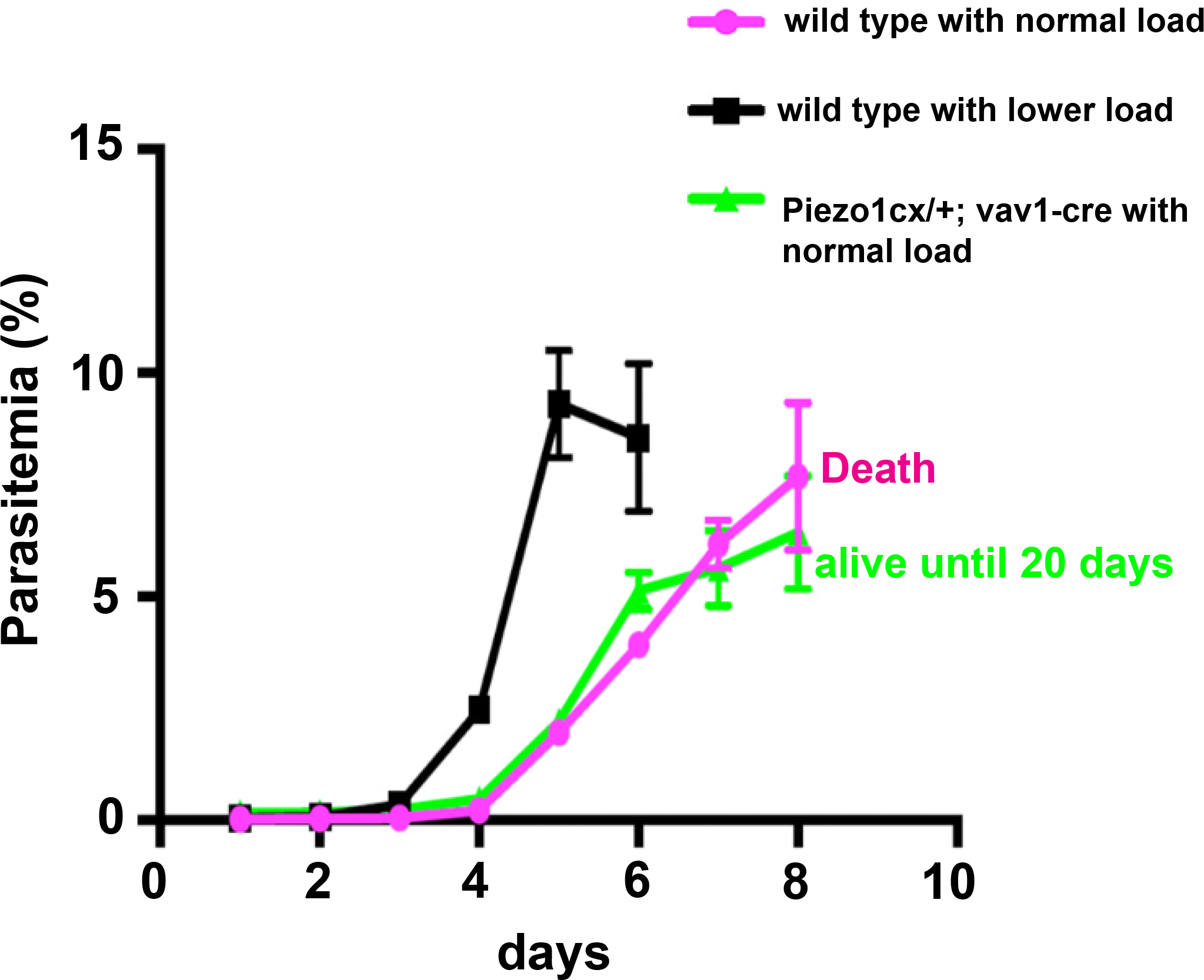
Wild type C57BL6 mice infected with a lower parasite load (4×10^4^) (pink color) had a similar parasitemia curve as blood specific Piezo1 GOF mice infected with 1×10^5^ iRBC (green) but died at day 8 in contrast to the Piezo1 GOF mice that survived until day 20.

**Figure S3.**
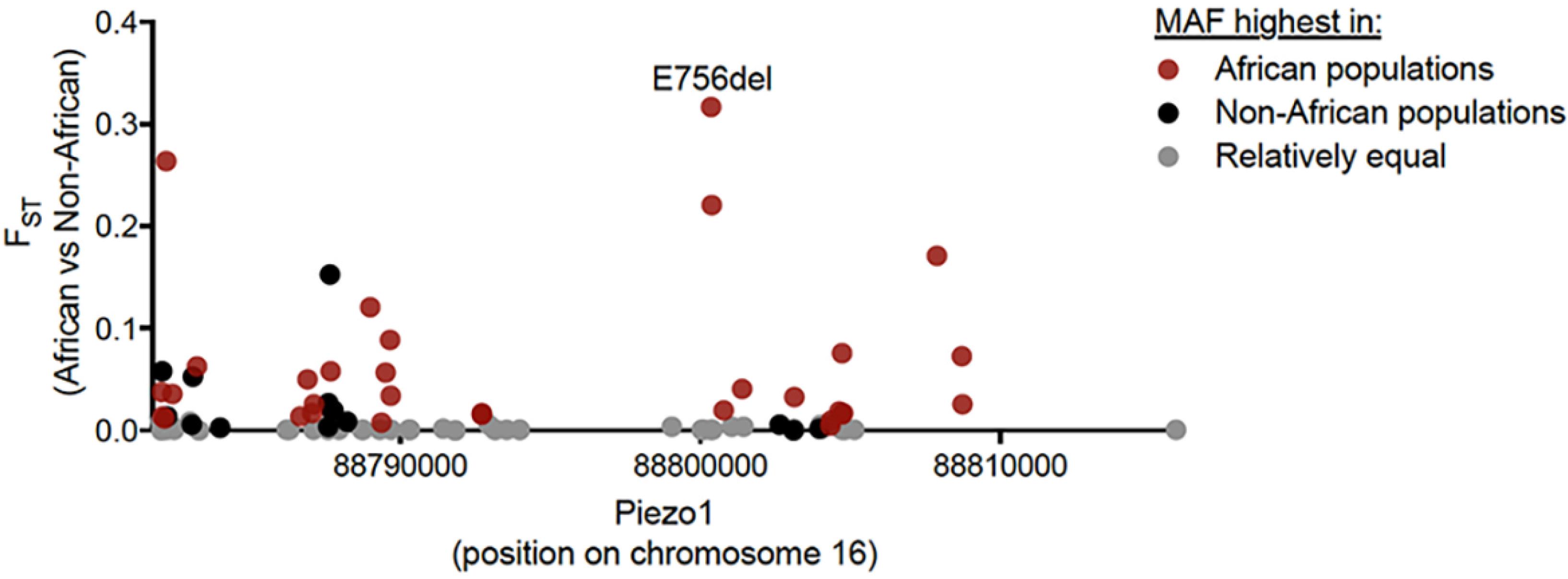
F_st_ values for Piezo1 missense and in-frame indel mutations. x-axis represents chromosome 16 coordinates. Red dots are alleles with MAF highest in African population while black dots are for non-Africans. Gray spots are alleles with similar or equal MAF in both populations.

**Supplemental Table 1.**
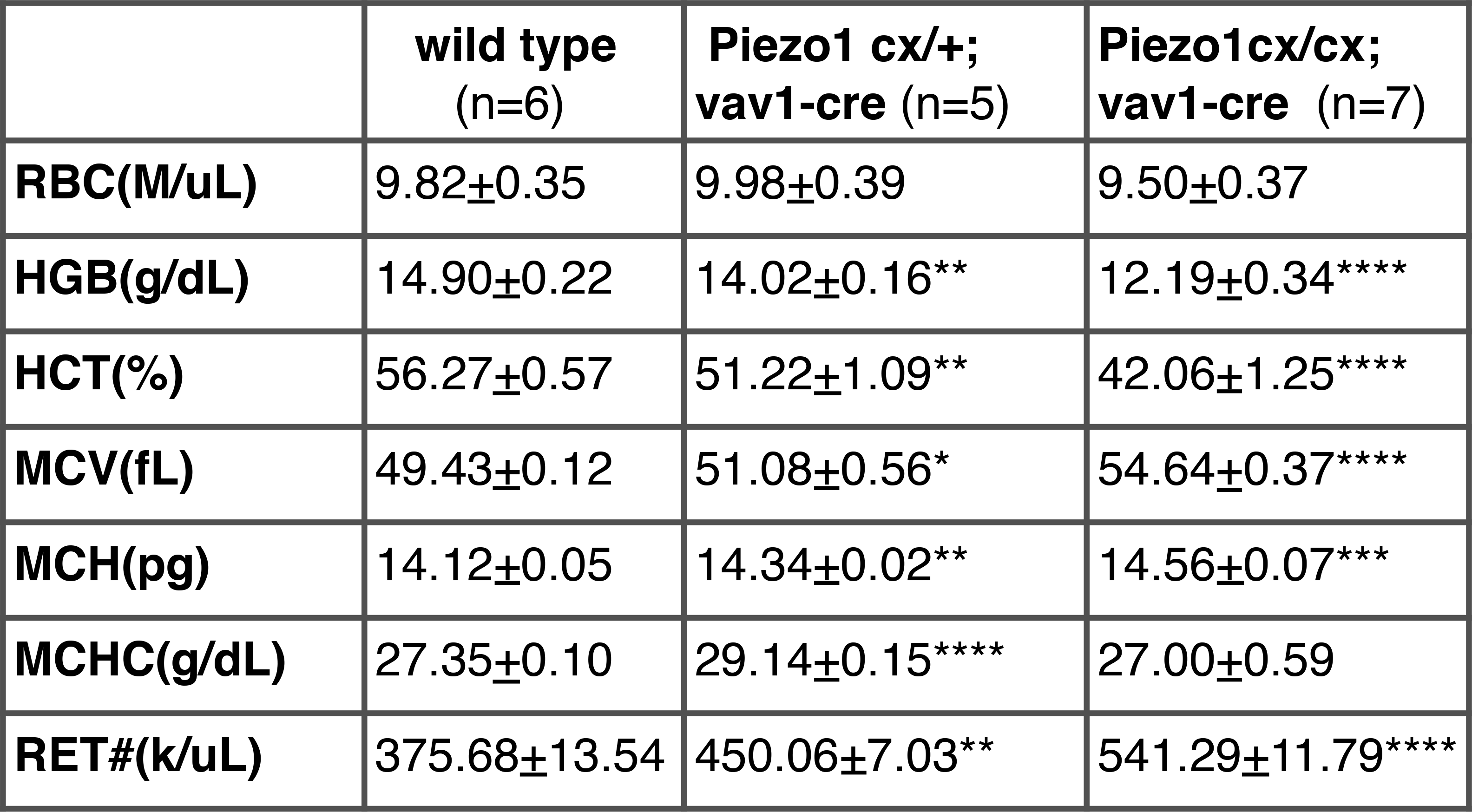
Complete cell counts on wildtype and Piezo1 cx/cx; vav1-cre mice. RBC- red blood cell. HGB- hemoglobin level. HCT- hematocrit. MCV- mean corpuscular volume. MCH- mean corpuscular hemoglobin. MCHC- mean corpuscular hemoglobin concentration. RET# - reticulocyte number (k=1000). Statistics: Student’s t test (between each knock-in line and wild type group). *p <0.05, **p <0.05, ***p <0.001, **** p <0.0001.

|  | **wild type**<br>(n=6) | **Piezo1 cx/+;**<br>**vav1-cre** (n=5) | **Piezo1cx/cx;**<br>**vav1-cre** (n=7) |
| --- | --- | --- | --- |
| **RBC(M/uL)** | 9.82±0.35 | 9.98±0.39 | 9.50±0.37 |
| **HGB(g/dL)** | 14.90±0.22 | 14.02±0.16** | 12.19±0.34**** |
| **HCT(%)** | 56.27±0.57 | 51.22±1.09** | 42.06±1.25**** |
| **MCV(fL)** | 49.43±0.12 | 51.08±0.56* | 54.64±0.37**** |
| **MCH(pg)** | 14.12±0.05 | 14.34±0.02** | 14.56±0.07*** |
| **MCHC(g/dL)** | 27.35±0.10 | 29.14±0.15**** | 27.00±0.59 |
| **RET#(k/uL)** | 375.68±13.54 | 450.06±7.03** | 541.29±11.79**** |

**Supplemental Table 2.**
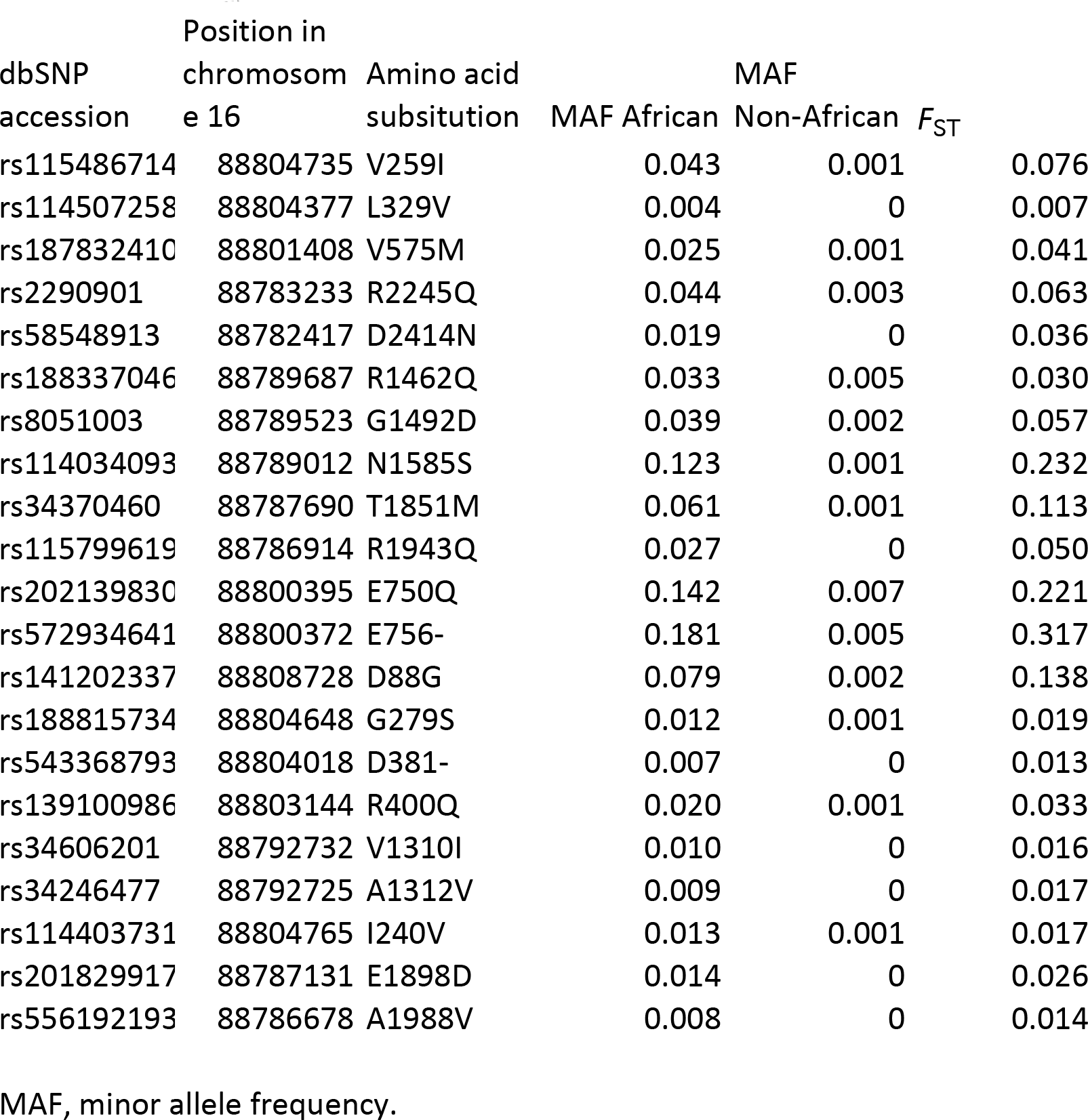
Piezo1 missense and in-frame indel mutations enriched in African population.

## Acknowledgements

We thank Ali Torkamani for advice on human genomics, Dominic Kwiatkowski for discussions, and Lisa Stowers for reading the manuscript. This work was partly supported by the following grants: NIH R01 DE022358 to A.P and AI090141 and AI103058 to EAW. S.M is supported by Calibr-GHDDI Gates postdoctoral fellowship. GLM is supported by an A.P. Giannini postdoctoral fellowship. A.P. is an investigator of the Howard Hughes Medical Institute. The authors declare no competing financial interests.

## Method

### Mouse lines

All animal procedures were approved by the Institutional Animal Care and Use Committees of The Scripps Research Institute (TSRI).

Piezo1cx/cx; vav1-cre, Piezo1cx/+; vav1-cre, Piezo1cx/cx; cmv-cre, Piezo1cx/+; cmv-cre mice were generated by breeding Piezo1 cx/cx with vav1-cre (12) (The Jackson Laboratory, stock# 018968) and cmv-cre(16) (The Jackson Laboratory stock# 006054), respectively. Piezo1 GOF mice were generated and maintained on C57BL/6 background. Both vav1-cre and cmv-cre mice were backcrossed at least 10 generations to C57BL/6.

### *P. berghei* infections and parasitemia measurement by flow cytometry

Donor mice (C57BL/6) were intraperitoneally injected with 50-200μl of erythrocytes parasitized with *P. berghei* (ANKA) GFPcon 259cl2. Blood was collected by cardiac puncture from infected donors when the parasitemia reached 4-6% (see below). Parasitized erythrocytes were washed with sterile saline three times at 1000xg for 3min and diluted to 5×10^5^ infected cells/ml as working solution. 200μl working solution was intravenously injected into the experimental mice for analysis. For GFP-fluorescence based parasitemia measurement, 1.5μl tail blood was collected from infected mice in 180μl Dulbecco’s Phosphate Buffer Solution with 2% fetal bovine serum, on 96-well plates. The cytometry was performed on NovoCyte Flow Cytometer system (ACEA Biosciences, San Diego, CA) following manufacturer’s instructions. Briefly, erythrocytes were selected on size for analysis by gating on forward/side-light scatter. Excitation of erythrocytes was performed with a laser at a wavelength of 488 nm and emission of the green fluorescence was detected using a filter of 530 nm. By gating the uninfected erythrocytes and the GFP-positive infected erythrocytes parasitemia was calculated as the percentage of infected cells.

### Blood-brain barrier assay

When *P. berghei* infected displayed either experimental cerebral malaria (including ataxia, convulsion, paralysis and/or coma) or severe anemia (immobility and pale blood color), 2% Evans Blue (Sigma-Aldrich, dissolved in sterile saline) was intravenously injected into *P. berghei* infected at 5ml/kg body weight. After 45-60 min, euthanized mice were transcardially perfused with Phosphate Buffer Solution and 4% paraformaldehyde before brains were dissected.

### Scanning Electron Microscope

Samples of mouse blood were added to ice cold buffered saline (10mM NaCl, 155mM KCl, 10mM glucose and 1 mM magnesium chloride) before being fixed in 2.5% glutaraldehyde in 0.1 M cacodylate buffer on ice. Aliquots of the fixed cells were placed on 12mm coverslips previously coated in polylysine for 30 mins. Following a buffer wash and postfixation in buffered 1% osmium tetroxide, the samples were extensively washed in distilled water. The samples were dehydrated in graded ethanol series followed by processing in a critical point drier (Tousimis autosamdri 815). The coverslips were then mounted onto SEM stubs with carbon tape and sputter coated with Iridium (EMS model 150T S) for subsequent examination and documentation on a Hitachi S-4800 SEM (Hitachi High Technologies America Inc., Pleasanton CA) at 5kV.

For human RBCs, samples were fixed at room temperature with 2.5% glutaraldehyde in 0.1M cacodylate buffer followed by 1% osmium tetroxide, and washed in water. A 10 µl drop of suspension was loaded on the sample carrier and imaged in a FEI Quanta200 FEG microscope in ESEM mode using the gaseous secondary electron detector. The stage was set-up at 2°C, the acceleration voltage was 15kV and the working distance 10 mm. Water was then progressively removed by cycles of decreasing pressure / injection of water, until reaching equilibrium at the dew point. The minimal final pressure in the chamber was 350 Pa. Pictures were taken with a dwell time of 6 μsec.

### Osmotic fragility test

Blood was diluted at 1:8 into isotonic saline (0.9% NaCl) containing 2 mM HEPES, pH 7.4. 10 µl of the diluted blood was pipetted into each well (in a row) on a 96-well round-bottom plate. Separate rows were used for separate samples. 225 μl tonicity solutions made from saline solutions at concentration of 0, 20, 25, 30, 35, 40, 45, 50, 55, 60, 80, and 100%. The plate was incubated for 5min at room temperature followed by 5 min centrifuge at 1000xg. 150 µl supernatant was transferred to 96-well flat bottom plate for absorbance reading at 540nm using Cytation3 (BioTek, Winooski, VT). The data was analyzed using 4-parameter sigmoidal nonlinear regression.

### Piezol GOF mice generation

The targeting strategy was based on the NCBI transcript NM_001037298.1. Wildtype exons 45-51, including the complete 3’ untranslated region (UTR) were flanked by loxP sites. An additional polyadenylation signal (nucleotide sequence of the synthetic polyA: gagctccctggcgaattcggtaccaataaaagagctttattttcatgatctgtgtgttggtttttgtgtgcggcgcg) was inserted between the 3’ UTR and the distal loxP site in order to prevent downstream transcription of the mutated exon 51 in the conditional allele. The size of the loxP-flanked region is 2.8 kb. The exons 45-51, including the splice acceptor site of intron 44 were duplicated and inserted downstream of the distal loxP site. The R2482H mutation was introduced into the duplicated exon 51. Positive selection markers were flanked by FRT (Neomycin resistance - NeoR) and F3 (Puromycin resistance-PuroR) sites and inserted into intron 44 and downstream of the synthetic polyA, respectively. The targeting vector was generated using BAC clones from the C57BL/6J RPCIB-731 BAC library and transfected into the TaconicArtemis C57BL/6N Tac ES cell line (Taconic, Hudson, NY). Homologous recombinant clones were isolated using double positive (NeoR and PuroR) selection. The conditional knock-in allele was obtained after Flp-mediated removal of the selection markers. The constitutive knock-in allele was obtained after Cre-mediated deletion of wildtype exons 45-51 and the synthetic polyA sequences

### Mechanical stimulation

For whole-cell recordings, mechanical stimulation was achieved using a fire-polished glass pipette (tip diameter 3–4 μm) positioned at an angle of 80° to the recorded cells. Downward movement of the probe towards the cell was driven by a Clampex controlled piezo-electric crystal microstage (E625 LVPZT Controller/Amplifier; PhysikInstrumente). The probe had a velocity of 1 μm.ms^-1^ during the ramp segment of the command for forward motion and the stimulus was applied for 150 ms. To assess the mechanical responses of a cell, the probe was first placed as close to the cell as possible (this distance could vary from cell to cell). Then, a series of mechanical steps in 1 μm increments was applied every 10s, which allowed full recovery of mechanically activated (MA) currents. Threshold was calculated by subtracting the distance at which the probe first touched the cell surface from the minimal distance at which mechanically activated currents were evoked. Mechanically activated inward currents were recorded at a holding potential of −80 mV. The inactivation kinetics at a holding potential of −80 mV of traces of currents reaching at least 75 % of the maximal amplitude of current elicited per cell were fitted with mono-exponential equation (or in some case bi-exponential equation for the rapidly-adapting currents, accordingly to previous reports (6) and using the fast time constant giving a value of inactivation time (τ) per responsive cell used for analysis. Channel kinetic properties between WT and mutant Piezo1 were compared using Student’s t-test.

### Cell culture and transient transfection

Piezo1 KO HEK cells (11) were grown in Dulbecco’s modified Eagle’s medium containing 4.5 mg/ml glucose, 10% fetal bovine serum, 1 × antibiotics/antimycotics. Cells were plated in 6-well plates and transfected using lipofectamines 2000 (Invitrogen by ThermoFisher Scientific, Carlsbad, CA), according to the manufacturer’s instruction. Human or mouse Piezo1 mutations fused to IRES-tdTomato was transfected at 1.4μg per well (6-well plate) for Fluorescent imaging plate reader (see below). To measure calcium signals, ultrasensitive sensor GCaMP6 (29) were transfected at 0.6 µg per well (6-well plate). Cells were incubated for 2 days before electrophysiology experiments.

### Fluorescent imaging plate reader (384-well format)

After transfection, the cells were dissociated from 6-well plates and re-seeded into 384-well plate, at 12,000 cells per well. The plate was incubated for 2 days then washed with assay buffer (1 × HBSS, 10 mM HEPES, pH7.4) using a ELx405 CW plate washer (BioTek, Winooski, VT). Fluorescence was monitored on a fluorescent imaging plate reader (FLIPR) Tetra. A 10-mM stock solution of Yoda1 in dimethyl sulfoxide (DMSO) was used resulting in a maximum of 1% DMSO in the assay. 10 µM Yoda1 was used in initial screens for searching gain-of-function mutations (compared to wild type). Positive hits were then validated by using a series of Yoda1 concentrations. Concentration-response curves were fitted using a sigmoidal dose-response at variable slope (GraphPad Prism, La Jolla, CA).

### Real time quantitative PCR

Total RNA was isolated from mouse whole blood by Quick-RNA Whole Blood (Zymo Research, Irvine, CA). 500 ng total RNA was used to generate 1st strand cDNA using the Quantitect reverse transcription kit (Qiagen). Real time PCR assays were set up using GoTaq qPCR Master Mix (Promega, Madison WI). The reaction was run in the ABI 7900HT fast real time system using 1 μl of the cDNA in a 20 μl reaction according to the manufacturer’s instructions in triplicates. Primers were designed for target gene (mPiezo1) (forward CTCACAGACAGGTGTTCATC, reverse GCAAACTCACGTCAAGGAGA) and reference gene (Gapdh) (forward primer GCACCACCAACTGCTTAG, reverse primer GGATGCAGGGATGATGTTC). Calibrations and normalizations were done using the 2-ΔΔCT method, where ΔΔCT = ((CT(target gene) -CT (reference gene)) - (CT (calibrator) - CT (reference gene)).

### Population genetic analysis

We obtained minor allele and genotype frequencies from the Exome Aggregation Consortium (ExAC) (19) and the 1000 Genomes Project (21). 2504 genomes were analyzed, 661 from African and 1843 from non-African ancestries. Wright’s fixation index (F_ST_), a measure of population differentiation, was calculated as follows:

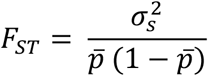

where 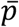 is the mean allele frequency and 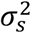 is the allele frequency variance between the populations. The 1000 Genomes browser (http://phase3browser.1000genomes.org) was used to determine that no alleles were in linkage disequilibrium with E756del (estimated r^2^ values were <0.05). Hardy-Weinberg equilibrium was estimated using the classical binomial expansion to determine the expected genotype frequencies and x^2^ tests.

### Yoda1 treatment on human RBCs for *P. falciparum* infection

For Yoda1 treatment of *P. falciparum* cultures, 3D7 strain (MRA-102 from MR4) was cultured in human RBC (obtained from the local blood bank) at 5% hematocrit in complete medium composed of RPMI 1640 medium supplemented with 25 mM HEPES buffer (pH 7.4), 0.5% albumax I (Thermo Fisher Scientific), and gentamycin (40 μg/ml, Gibco). Cultures were incubated shaking at 37°C in a culture gas chamber under a gaseous mixture of 5% CO_2_, 5% O_2_ and 90% N_2_. For experiments, late stage parasites from synchronous cultures were purified by passage over VarioMACS columns (Miltenyi Biotec), washed and eluted with complete medium and treated with 1.5 μM of the protein kinase G inhibitor compound2 for up to 5 h. This allows parasites to develop to segmenters but blocks egress until inhibitor is removed by washing. These preparations were then mixed with either Yoda1 pre-treated RBCs or with untreated RBCs and Yoda1 added just after mixing. To treat ring stage parasites, purified parasites were mixed with RBCs, incubated for 4 h and synchronized by 5% sorbitol treatment. Cultures were allowed to stabilize for 1 h before adding Yoda1. For determination of parasitemia by FACS analysis, cultures were centrifuged and the cells fixed for at least 4 h at room temperature in 10 volumes of 4% paraformaldehyde (Electron Microscopy Sciences) in phosphate buffered saline (PBS). Preparations were then washed twice with PBS and incubated 30 min with 3.3×SYBR Green (Thermo Fisher Scientific). DNA dye was removed by a simple wash in PBS and samples analyzed on a FACS Canto (BD Biosciences) using processed uninfected RBCs as control.

### *P. falciparum* culture

*P. falciparum* Dd2 strain parasites were cultured under standard conditions (30), using RPMI media supplemented with 0.05 mg/ml gentamycin, 0.014 mg/ml hypoxanthine (prepared fresh), 38.4 mM HEPES, 0.2% Sodium Bicarbonate, 3.4 mM Sodium Hydroxide, 0.05% O+ Human Serum (Denatured at 56°C for 40 min; Interstate Blood Bank, Memphis, TN) and 0.0025% Albumax). Human O+ whole blood was obtained from TSRI Normal blood donor service (La Jolla, CA). Leukocyte-free erythrocytes are stored at 50% hematocrit in RPMI-1640 screening media (as above, but without O+ human serum and with 2× albumax concentration) at 4°C for one to three weeks before experimental use. Cultures were monitored every one to two days via Giemsa-stained thin smears.

### Parasitemia Determination

Asynchronous *P. falciparum* parasites (Dd2 strain) were cultured in standard conditions (as described above), then synchronized twice via sorbitol (31) and grown to 7% parasitemia at the late trophozoite/schizont stage. Patient blood was obtained from TSRI Normal blood donor service, washed and centrifuged three times (at 800 × g for 5 min at 4°C) with RPMI only and once with complete media (as above) (32), with any visible buffy coat being removed after each spin. All blood samples were given a numerical designation and allele status was not determined until after all data collection was completed.

Parasite growth was determined in two independent ways: absolute parasitemia determination via thin-blood smear after Giemsa staining, and inferred parasite growth via the SYBR Green I-based fluorescence assay (33). For the absolute parasitemia determination by thin blood smear, parasites were established, starting from the 7% parasitemia cultures above and diluted using the corresponding patient RBCs, in a 10 mL culture at 5% hematocrit and 0.5% parasitemia (the same as for the Giemsa absolute parasitemia determination assay). Cultures were then grown for 4 days and parasitemia measurements were taken every 2 days. For the SYBR green I inferred parasitemia determination assay, when parasite burden is estimated based upon DNA content, parasites were cultured in 100 μL volumes in 96-well plates at 5% hematocrit and 0.5% parasitemia with at least 5 replicates per time point. Parasites were plated on three 96-well black plates with clear bottoms (Fisher Scientific). (One surrogate plate was used for measurement every two days, with DNA content determined by SYBR Green I incorporation of lysed parasites). Relative parasitemia was determined by fluorescence measurement, background was determined using uninfected RBCs and subtracted, then relative parasite burden was determined via normalization against a known O+ WT blood sample. In both cases all measurements were taken for all samples, genotypes were then assigned to numbered patient samples, wild type vs heterozygote samples were averaged at each time point, and average parasitemia values were compared.

## References

1. D.P. Kwiatkowski, How malaria has affected the human genome and what human genetics can teach us about malaria. Am. J. Hum. Genet. 77, 171–192 (2005).

2. Z. Feng, D.L. Smith, F.E. McKenzie, S.A. Levin, Coupling ecology and evolution: malaria and the S-gene across time scales. Math Biosci. 189, 1–19 (2004).

3. P. Hedrick, Estimation of relative fitnesses from relative risk data and the predicted future of haemoglobin alleles S and C. J. Evol. Biol. 17, 221–224 (2004).

4. E. Elguero et al., Malaria continues to select for sickle cell trait in Central Africa. Proc. Natl. Acad. Sci. U.S.A. 112, 7051–7054 (2015).

5. J. Delaunay, The hereditary stomatocytoses: genetic disorders of the red cell membrane permeability to monovalent cations. Semin. Hematol. 41, 165–172 (2004).

6. J. Albuisson et al., Dehydrated hereditary stomatocytosis linked to gain-of-function mutations in mechanically activated PIEZO1 ion channels. Nat Commun. 4, 1884 (2013).

7. R. Zarychanski et al., Mutations in the mechanotransduction protein PIEZO1 are associated with hereditary xerocytosis. Blood. 120, 1908–1915 (2012).

8. C. Bae, R. Gnanasambandam, C. Nicolai, F. Sachs, P.A. Gottlieb, Xerocytosis is caused by mutations that alter the kinetics of the mechanosensitive channel PIEZO1. Proc. Natl. Acad. Sci. U.SA. 110, E1162–1168 (2013).

9. S. M. Cahalan et al., Piezo1 links mechanical forces to red blood cell volume. Elife. 4 (2015), doi:10.7554/eLife.07370.

10. T. Tiffert et al., The hydration state of human red blood cells and their susceptibility to invasion by Plasmodium falciparum. Blood. 105, 4853–4860 (2005).

11. A. E. Dubin et al., Endogenous Piezo1 Can Confound Mechanically Activated Channel Identification and Characterization. Neuron. 94, 266–270.e3 (2017).

12. J. de Boer et al., Transgenic mice with hematopoietic and lymphoid specific expression of Cre. Eur. J. Immunol. 33, 314–325 (2003).

13. N. M. Archer et al., Hereditary xerocytosis revisited: Hereditary Xerocytosis Revisited. American Journal of Hematology. 89, 1142–1146 (2014).

14. B. Franke-Fayard et al.. A Plasmodium berghei reference line that constitutively expresses GFP at a high level throughout the complete life cycle. Mol. Biochem. Parasitol. 137, 23–33 (2004).

15. M.M. de Oca, C. Engwerda, A. Haque Plasmodium berghei ANKA (PbA) infection of C57BL/6J mice: a model of severe malaria. Methods Mol. Biol. 1031, 203–213 (2013).

16. F. Schwenk, U. Baron, K. Rajewsky A cre-transgenic mouse strain for the ubiquitous deletion of loxP-flanked gene segments including deletion in germ cells. Nucleic Acids Res. 23, 5080–5081 (1995).

17. R. Idro, N.E. Jenkins, C. R. J. C. Newton, Pathogenesis, clinical features, and neurological outcome of cerebral malaria. Lancet Neurol. 4, 827–840 (2005).

18. R.E. Phillips, G. Pasvol, Anaemia of Plasmodium falciparum malaria. Baillieres Clin. Haematol. 5, 315–330 (1992).

19. R. Syeda et al.. Chemical activation of the mechanotransduction channel Piezo1. Elife. 4 (2015), doi:10.7554/eLife.07369.

20. M. Lek et al.. Exome Aggregation Consortium, Analysis of protein-coding genetic variation in 60,706 humans. Nature. 536, 285–291 (2016).

21. A. Auton et al. with 1000 Genomes Project Consortium, A global reference for human genetic variation. Nature. 526, 68–74 (2015).

22. J. J. Vitti, S.R. Grossman, P.C. Sabeti, Detecting natural selection in genomic data. Annu. Rev. Genet. 47, 97–120 (2013).

23. Basic Malaria Microscopy. Part I. Learner’s Guide, Second Edition. WHO. 2010.

24. Malaria Genomic Epidemiology Network, Malaria Genomic Epidemiology Network, Reappraisal of known malaria resistance loci in a large multicenter study. Nat. Genet. 46, 1197–1204 (2014).

25. E.M, Leffler et al.. Malaria Genomic Epidemiology Network, Resistance to malaria through structural variation of red blood cell invasion receptors. Science. 356 (2017), doi:10.1126/science.aam6393.

26. S. S. Ranade et al., Piezo1, a mechanically activated ion channel, is required for vascular development in mice. Proc. Natl. Acad. Sci. U.S.A. 111, 10347–10352 (2014).

27. J. Li et al.. Piezo1 integration of vascular architecture with physiological force. Nature. 515, 279–282 (2014).

28. S. Wang et al.. Endothelial cation channel PIEZO1 controls blood pressure by mediating flow-induced ATP release. J. Clin. Invest. 126, 4527–4536 (2016).

29. T.-W. Chen et al., Ultrasensitive fluorescent proteins for imaging neuronal activity. Nature. 499, 295–300 (2013).

30. W. Trager, J.B. Jensen, Human malaria parasites in continuous culture. Science. 193, 673–675 (1976).

31. C. Lambros, J.P. Vanderberg, Synchronization of Plasmodium falciparum erythrocytic stages in culture. J. Parasitol. 65, 418–420 (1979).

32. J.M. Elias, C. Greene, Modified Steiner method for the demonstration of spirochetes in tissue. Am. J. Clin. Pathol. 71, 109–111 (1979).

33. J.D, Johnson et al., Assessment and continued validation of the malaria SYBR green I-based fluorescence assay for use in malaria drug screening. Antimicrob. Agents Chemother. 51, 1926–1933 (2007).

